# The Role of Vitamin D in *Emiliania huxleyi*: A Microalgal Perspective on UV Exposure

**DOI:** 10.1101/2023.09.21.558789

**Authors:** Or Eliason, Sergey Malitsky, Irina Panizel, Ester Feldmesser, Ziv Porat, Martin Sperfeld, Einat Segev

**Affiliations:** Department of Plant and Environmental Sciences, Weizmann Institute of Science; Rehovot, 7610001, Israel; Department of Life Sciences Core Facilities, Weizmann Institute of Science; Rehovot, 7610001, Israel

## Abstract

An essential interaction between sunlight and eukaryotes involves the production of vitamin D through exposure to ultraviolet (UV) radiation. While extensively studied in vertebrates, the role of vitamin D in non-animal eukaryotes like microalgae remains unclear. To investigate the potential involvement of vitamin D in the response of microalgae to UV, we focus on *Emiliania huxleyi*, a microalga found in shallow ocean depths that are exposed to UV radiation. Our results show that *E. huxleyi* algae produce vitamin D_2_ and D_3_ in response to UV irradiation. We further demonstrate that *E. huxleyi* algae respond to external administration of vitamin D at the transcriptional level, regulating the expression of protective mechanisms that are also regulated in response to UV. Our data reveal that addition of vitamin D enhances the algal photosynthetic performance while reducing harmful reactive oxygen species buildup. This study contributes to understanding the function of vitamin D in *E. huxleyi* and sheds light on its role in non-animal eukaryotes, as well as its potential importance in marine ecosystems.

## Introduction

Life on Earth has a complex relationship with sunlight, relying on its energy for certain processes while simultaneously requiring protection against its potential harmful effects. A molecular process that is tightly linked to sunlight is the formation of vitamin D following exposure to ultraviolet-B (UV-B) radiation emitted from the sun. Vitamin D (calciferol) comprises a group of steroids that result from the photochemical transformation of several sterol precursors by UV-B wavelengths^1^. The most common vitamin D species known to occur naturally are vitamin D_2_ and D_3_, originating from the conversion of ergosterol and 7-dehydrocholesterol, respectively^2,3^.

In mammals and other studied vertebrates, vitamin D functions as a hormone, involved in the regulation of a multitude of intracellular and physiological processes vital for the organism survival and well-being^4^. Vitamin D is pivotal in facilitating the absorption and homeostasis of essential ions such as Ca^2+^ and PO_4_^3–5,6^, and vitamin D deficiency has been linked to a range of physiological disorders^7,8^. Due to its key role in human health, vitamin D has been the focus of biological and pharmaceutical research efforts, directed towards understanding the mechanisms through which it operates in humans and human models.

However, mounting evidence suggest that vitamin D has been a constituent of eukaryotes long before the emergence of vertebrates. This is evident not only in the identification of vitamin D in distant eukaryotic lineages like algae^9–11^, plants^12^, and fungi^13–15^ but also in the preservation of vitamin D-related biomarkers, likely from an algal source, in marine sediments dating back over 600 million years^16^.

Despite its widespread presence across diverse lineages, our understanding of the role of vitamin D in non-animal eukaryotes remains limited. Non-animal eukaryotes, namely microalgae, have been suggested as potential sources of vitamin D for higher trophic levels in the marine environment^17–19^. But the processes underlying vitamin D formation and regulation in microalgae remain largely unexplored^20^.

UV-B radiation is crucial for vertebrate health due to its role in vitamin D formation, but it can also be detrimental. UV can cause direct damage to biomolecules like DNA, leading to the generation of reactive oxygen species (ROS)^21–23^, and can ultimately result in cell death^24^. Photosynthetic organisms, like algae, are particularly susceptible to UV damage, as their energy production hinges on exposure to solar radiation^25,26^. Although water acts as a UV-B filter^27^, significant intensities can still penetrate the upper layers of the ocean^28^, potentially impacting organisms such as algae^29^.

Microalgae of the species *Emiliania huxleyi*, also named *Gephyrocapsa huxleyi*^30^, are widely distributed in modern oceans and play key roles in various biogeochemical cycles^31,32^. These algae are known to flourish in high light environments at shallow depths of about 10 to 20 meters^33^, where exposure to UV wavelengths is likely. Earlier findings provided intermittent indications that *E. huxleyi* algae might synthesize vitamin D. These reports highlighted the algal capacity to generate vitamin D_2_ upon exposure to UV-B irradiation^11^, the algal cholesterol content^34,35^, and the presence of a gene analogous to 7-dehydrocholesterol reductase (DHCR7) responsible for converting 7-dehydrocholesterol into cholesterol^36^. However, a comprehensive understanding of the cellular role of vitamin D in *E. huxleyi*, an environmentally key microalga, is still lacking.

In this study we explore the overlooked role of vitamin D in *E. huxleyi*. Specifically, we investigate vitamin D formation following exposure to UV and the regulation of cellular mechanisms that operate in response to harmful radiation.

## Results

### *E. huxleyi* algae produce vitamin D_2_ and D_3_

To investigate whether vitamin D is formed by *E. huxleyi* algae upon exposure to UV under our experimental settings, we cultivated algal cultures in a chamber with environmentally relevant UV-B radiation levels (see Materials and Methods). Metabolic analyses revealed the presence of both D_2_ and D_3_ in irradiated algal cultures (Table 1). Our results show that D_2_ was significantly enriched in UV-exposed cultures, with levels of approximately ∼4 ng/mg dry weight, while it was barely detected in cultures that were not exposed to UV. The D_2_ precursor ergosterol was found in both UV-treated and control cultures. Lower amounts of vitamin D_3_ (∼0.04 ng/mg dry weight) were detected in both UV-treated and control cultures.

**Table 1.** *E. huxleyi* algae produce vitamin D_3_ and D_2_. Metabolic analysis of vitamin D species and precursors under UV and control conditions, using algal cultures at day 10 of growth. Values are averages of dry weight (ng / mg) ± standard deviation based on 4 biological replicates. Statistically significant differences (*p <* 0.05) between treatments are marked by *, calculated using two-sample t-test assuming equal variances.

|  | UV | Control |
| --- | --- | --- |
| Compound | Dry weight (ng / mg) |  |
| D<sub>2</sub> | 4.32 ± 1.39 * | 0.09 ± 0.01 |
| D<sub>3</sub> | 0.038 ± 0.001 | 0.039 ± 0.001 |
| Ergosterol | 83.01 ± 28.96 | 71.76 ± 9.53 |
| 7-dehydrocholesterol | 0.24 ± 0.04 * | 0.16 ± 0.02 |

Importantly, while D_2_ detection was consistent across all analyzed UV-exposed samples, D_3_ was identified only in part of our experiments. When D_3_ was detected, its precursor 7-dehydrocholesterol was detected as well. Inconsistent detection of D_3_ was previously reported in plants and was attributed to the sensitivity of the analytical method used^37^. Our many efforts to resolve the variable measurements of D_3_ were not successful (see detailed description of attempts in Materials and Methods). Collectively, our findings demonstrate that *E. huxleyi* algae produce D_2_ and D_3_, with increased levels of D_2_ following UV irradiation. These observations suggest a possible role for vitamin D in the algal response towards UV.

### *E. huxleyi* algae show a transcriptomic response to UV radiation

UV radiation is necessary for the formation of vitamin D^2^. In other organisms, once vitamin D is generated, it functions as a hormone that regulates various physiological processes^4^. Therefore, we sought to explore the transcriptomic response of *E. huxleyi* algae to UV exposure, seeking to elucidate cellular processes that may be related to vitamin D. We therefore analyzed the *E. huxleyi* transcriptome in cultures that were grown under diurnal light conditions that included UV irradiation, in comparison to algal cultures that were protected from the UV source. Cultures were sampled for RNA sequencing at three time points representing different growth phases (days 7, 10, and 13, see Figure S1).

The transcriptomic analysis revealed differential expression of 374 genes between UV-exposed and control cultures (Table S1). Of these genes, 172 were annotated with GO terms related to a known function or process. The annotated genes that were differentially expressed in the transcriptome under UV exposure were associated with various cellular processes including intracellular signaling pathways and stress response mechanisms. Notably, genes participating in the inositol 3-phosphate/calcium (IP_3_/Ca^2+^) and the oxylipin signaling pathways were differentially expressed (Table 2), both playing key roles in stress response mechanisms across different organisms^38–42^.

**Table 2.** Differential expression (DE) of genes associated with signaling and stress response mechanisms in *E. huxleyi* that were upregulated under UV. DE values calculated according to the transcriptomic analysis are given for days 7, 10 and 13 of growth (designated d. 7, d. 10 and d. 13). Genes are ordered according to DE values at day 13. Full DE and NCBI accession data are presented in Table S1 and Data S1.

| Gene ID | Putative protein or domain | Differential expression |  |  | Protein function |
| --- | --- | --- | --- | --- | --- |
| Genes related to intracellular signaling |  | d. 7 | d. 10 | d. 13 |  |
| G10384 | Lipoxygenase domain | -0.46 | -1.27 | 2.04 | Oxylipin biosynthesis |
| G14992 | Prostaglandin F(2-alpha) synthase | 0.38 | 0.22 | 1.75 | Oxylipin biosynthesis |
| G12340 | Cytosolic phospholipase A2 domain | -0.51 | -0.82 | 1.70 | Oxylipin biosynthesis; Intracellular signaling |
| G14502 | Ca-binding domain | -0.57 | -1.12 | 1.67 | Shares similarity to *A. thaliana* calmodulin-like protein; Intracellular signaling. |
| G21784 | Phosphoinositide phospholipase C | -0.52 | -0.62 | 1.50 | IP<sub>3</sub> signaling initiation; cytosolic calcium regulation |
| G15496 | Phosphoinositide 5-phosphatase | 0.54 | 0.74 | 1.46 | IP<sub>3</sub> signaling; cytosolic calcium regulation |
| G25467 | Steroid hydroxylase | 1.15 | 1.49 | 1.16 | Shares similarity to rat *cyp1a2* with a suggested 25-cholesterol hydroxylase activity |
| G1648 | Mannosyl phosphorylinositol ceramide synthase | -0.37 | -0.21 | 0.98 | Potentially involved in Calcium signaling |
| G19702 | Phosphatidylinositol 3-kinase | -0.17 | 0.00 | 0.95 | IP<sub>3</sub> signaling |
| G27192 | Calcineurin B-like interacting protein kinase | -0.34 | 1.25 | -0.3 | Involved in Ca-mediated signaling; Abiotic stress response |
| Genes related to stress response |  |  |  |  |  |
| G27084 | Deoxyribodipyrimidine photolyase | 0.93 | 1.9 | 4.29 | UV-damage DNA repair |
| G647 | 3-dehydroquinate synthase domain | -0.31 | 5.33 | 4.07 | Shikimate pathway; potentially involved in the production of UV-protective compounds |
| G22973 | Sirtuin 2 | 1.16 | 0.89 | 1.49 | DNA transcription and repair |
| G16746 | Tyrosinase Cu-binding domain | -0.54 | -1.09 | 1.46 | Involved in synthesis of protective pigments and antioxidants |
| G18590 | Light-harvesting protein | 0.51 | 1.22 | 1.13 | Shares similarity to *C. reinhardtii* LHCSR; Alleviates photo-oxidative stress |
| G12503 | Poly ADP-ribose polymerase Zn-finger domain | 0.62 | 1.13 | 1.08 | DNA damage sensor |
| G18115 | Deoxyribodipyrimidine photolyase | 0.24 | 0.25 | 0.99 | UV-damage DNA repair |
| G6690 | Glutathione peroxidase | 0.05 | 0.32 | 0.98 | Oxidative stress mitigation |
| G25108 | Chalcone synthase 2 | -0.36 | -0.43 | 0.82 | Potentially involved in UV protection |
| G2876 | DNA-(apurinic or apyrimidinic site) lyase | 0.56 | 0.52 | 0.81 | DNA repair |
| G26038 | ATP-dependent DNA helicase | -0.34 | -0.76 | 0.79 | DNA stability and repair |
| G8907 | (6-4)DNA photolyase | 0.07 | 0.43 | 0.78 | UV-damage DNA repair |
| G2511 | Heme oxygenase | 0.37 | 0.96 | 0.67 | Oxidative stress mitigation |
| G16133 | Formamidopyrimidine-DNA glycosylase | 0.49 | 0.89 | 0.53 | DNA repair |
| G832 | Protochlorophyllide reductase | 1.23 | -0.41 | 0.2 | Chlorophyll biosynthesis |
| G22197 | 5'-tyrosyl DNA phosphodiesterase | 0.83 | 0.22 | 0.10 | DNA repair |

A substantial number of differentially expressed genes were involved in various stress responses, including DNA damage sensing and repair, oxidative stress mitigation, protective pigment biosynthesis, and maintenance of the photosynthetic machinery. Interestingly, several of the genes and pathways that were differentially expressed in *E. huxleyi* are known to be associated with UV exposure and vitamin D activity in vertebrates. For instance, the IP_3_/Ca^2+^ and oxylipin pathways are involved in UV stress response in mammals^43,44^, and vitamin D is involved in the regulation of these pathways^45–49^. In mammals, vitamin D also plays a role in oxidative stress mitigation, DNA repair, and the regulation of various enzymes related to stress responses including heme oxygenase, glutathione peroxidase, and tyrosinase^50–54^. Given that vitamin D regulates stress response mechanisms in mammals, and similar mechanisms are regulated by UV in *E. huxleyi*, vitamin D could potentially be involved in algal stress responses.

To explore the temporal dynamics of the algal response, specifically whether the algal transcriptomic response is elicited immediately upon exposure to UV, algal cultures were exposed to UV for a duration of 1 hour at day 10 of growth (Figures 2, S3). RNA was extracted and several genes that exhibited upregulation under diurnal UV conditions in our previous RNAseq analysis, were subsequently analyzed by qRT-PCR. The investigated genes showed significant upregulation. The differences in magnitude of differential expression observed between the two transcriptomic assays are possibly the result of a prolonged versus brief UV exposure. Taken together, algal cells exhibit a transcriptomic response following UV irradiation, including the activation of various stress response mechanisms. At least several of these mechanisms are immediately activated following a brief exposure to UV.

### External application of vitamin D impacts algal traits similarly to UV irradiation

To study the role of vitamin D in the algal response towards UV, we wished to manipulate the algal exposure to vitamin D. Exogenous application of vitamin D is a common practice in mammalian cells^53,55,56^. However, no knowledge exists on whether single-celled algae respond to this treatment. Therefore, to establish a vitamin D treatment in our algal cultures we screened a range of vitamin D concentrations and measured the effect on algal growth. An impact on algal growth would suggest that algal cells are affected by the exogenous addition of vitamin D.

Growth inhibition (*p* < 0.05) was observed under treatment of 1 µM of D_2_ and under the combination of 0.5 µM D_2_ and 0.5 µM D_3_ (Figure S2), suggesting that algae are affected by these treatments. The combination of D_2_ and D_3_ resulted in stronger growth inhibition compared to D_2_ alone (*p* = 0.00013). Importantly, under UV irradiation, algal growth inhibition was previously observed^57,58^, though our initial cultivation experiments did not demonstrate similar inhibition (Figure S1). We therefore adjusted our experimental setup to recapitulate the reported growth inhibition, and indeed observed a similar impact when the culture volume was reduced from 50 to 20 ml (Figure S2C), potentially due to shorter passage of UV in the medium.

We further investigated whether additional algal traits are similarly affected by UV irradiation and vitamin D treatment. Therefore, we examined cellular parameters that have been previously reported to be impacted by UV, including cell size, chloroplast size, and chlorophyll content^57,59^. Our findings show that when algae are cultivated under diurnal UV irradiation or are supplemented with a combination of D_2_ + D_3_ (0.5 µM of each), similar changes in algal traits are observed (Figures 1, S4). These changes include an increase in cell area, overall chloroplasts area, and chlorophyll a content. Furthermore, both treatments have resulted in a significant increase of the average number of chloroplasts per cell. An increase in chloroplast number, coupled with cell enlargement, is indicative of cell cycle arrest and is typical of UV-induced stress^57^. Additionally, a high correlation between cell and overall chloroplasts area was observed across biological replicates of control, vitamin D and UV treatments (Pearson correlation coefficient of r = 0.99, *p* = 4 * 10^-7^). These findings are in agreement with previous reports on UV effects in *E. huxleyi* and other microalgal species^57,59^. Importantly, other environmental stresses such as high light and nutrient starvation can also inhibit algal growth^60,61^. However, the impact of most environmental stresses on algal and plant physiology is reduction in chloroplasts to cell size ratio and in chlorophyll content^62,63^, contrary to the uniformity between treatments observed here (Figures 1, S4). Thus, both UV irradiation, and exogenous addition of D_2_ + D_3_ affect algal growth and several cellular traits in a similar manner.

**Figure 1.**
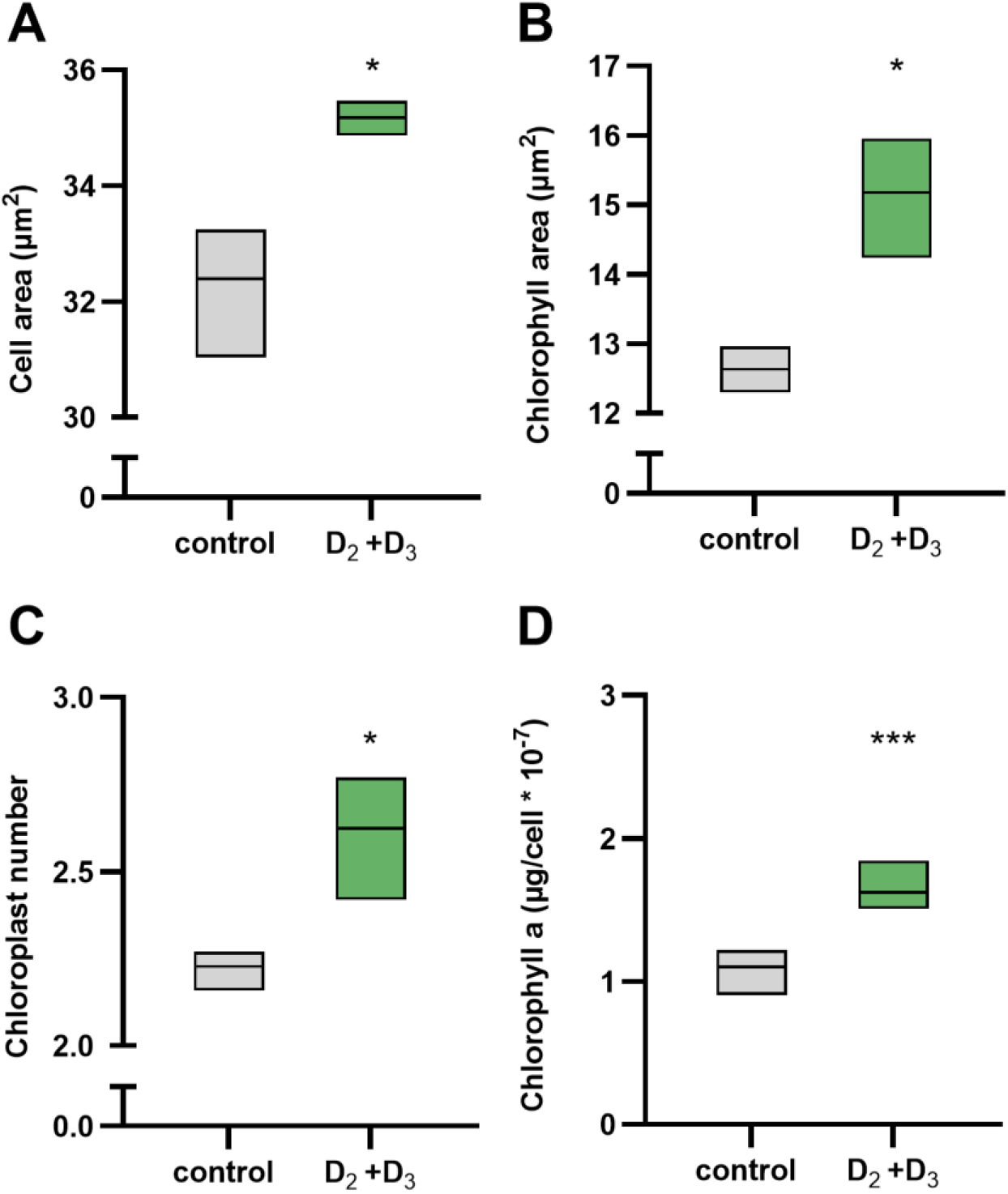
Vitamin D treatment leads to similar cellular traits as exposure to UV. (A) cell area, (B) chlorophyll area, (C) average number of chloroplasts and (D) cellular chlorophyll a content at day 10 of growth. Cultures were either treated with both D_2_ + D_3_ (0.5 µM of each) or DMSO (untreated) as control. Statistical significance of treated cultures compared to control conditions was calculated based on three biological replicates for (A-C) and six biological replicates for (d) using two-tailed t-test assuming equal variances. One, two or three asterisks indicate *p <* 0.05, *p* < 0.01 and *p <* 0.001, respectively.

**Figure 2.**
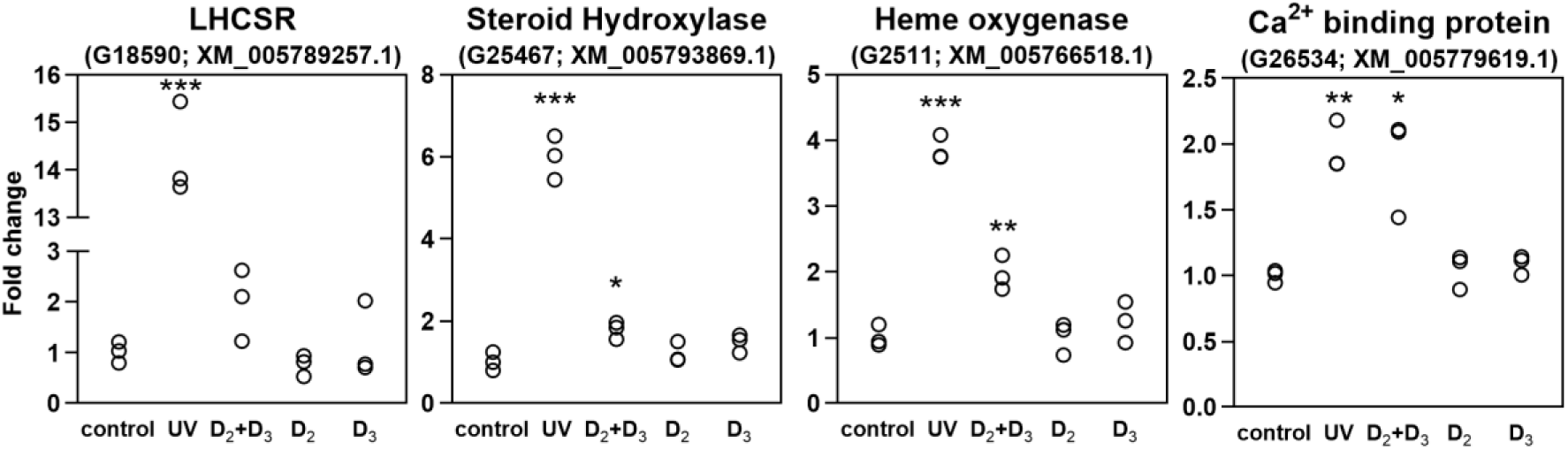
Combined treatment of vitamin D_2_ and D_3_ upregulates UV-responsive genes. qRT-PCR analysis of genes following 1 hour of UV exposure or vitamin D treatments. Top title denotes gene products. In brackets: gene identifier in *E. huxleyi* CCMP3266^94^ and matching gene transcript in the *E. huxleyi* CCMP1516 reference genome^95^. Upregulation of G18590 by vitamin D was observed for only two out of three replicates. Statistical significance of treated cultures compared to control conditions was calculated using two-tailed t-test assuming equal variances. One, two or three asterisks indicate *p <* 0.05, *p <* 0.01 and *p <* 0.001, respectively.

### Vitamin D upregulates expression of UV-regulated genes

After implementing a vitamin D treatment, we could examine whether vitamin D drives patterns of gene expression that are comparable those observed under UV irradiation. Similar expression patterns would suggest that vitamin D regulates a similar response to the response activated by UV. Therefore, we supplemented algal cultures with vitamin D and monitored the expression of UV-responsive genes via qRT-PCR.

Algal cultures were treated with 1 µM of D_2_ or D_3_, or with a combination of both (0.5 µM of each). Control cultures were supplemented with DMSO and exposed to either normal growth conditions or to UV radiation for 1 hour. RNA was collected from all cultures 1 hour post treatment.

Out of 12 monitored genes, our analyses revealed four genes that exhibit upregulated expression upon vitamin D addition and under UV irradiation (Figure 2). Notably, in vitamin D-treated cultures, the upregulation was only observed following addition of both D_2_ and D_3_. The four upregulated genes encode for light-harvesting complex stress-related protein (LHCSR, G18590), heme oxygenase (G2511), steroid hydroxylase (G25467), and a Ca-binding protein (G26534).

Considering the potential role of these four genes in a vitamin D-mediated response to UV, we searched the literature for their functions. Both LHCSR and heme oxygenase are proteins related to the algal stress response. The LHCSR gene is known to exhibit increased expression in moss and green algae under UV-B and high-light stress, promoting excess energy dissipation in the light-harvesting complex, thereby reducing photo-oxidative stress^64–66^. Heme oxygenases are enzymes involved in the formation of antioxidants in plants and animals, with known upregulated expression in response to UV-B and other ROS-forming stressors^67–69^. Notably, UV-B radiation is a fundamental component of high light environments. As vitamin D is produced under UV radiation (Table 1) and triggers the upregulation of oxidative and photooxidative stress mitigation pathways, it is possible that vitamin D is part of a cellular response towards harmful light intensities or radiation.

### Vitamin D treatment improves the algal photosynthetic performance following exposure to saturating light

To investigate the involvement of vitamin D in the algal response towards harmful light intensities, we tested the impact of D_2_ + D_3_ (0.5 µM of each) on algal photosynthetic performance under excess light. Algae were subjected to two light regimes: first, following a 5-minute dark-incubation period, algal cultures were exposed to saturating light intensities (1150 μmol photons m^−2^ s^−1^) and their photosynthetic parameters were evaluated. Additionally, we subjected algae to fluctuating light, common to the marine environment^70^. To this end, algal cultures were initially exposed to high light intensities (1000 μmol photons m^−2^ s^−1^) for 2 hours before undergoing a 5-minute dark-incubation period, followed by a second exposure to high light. Photosynthetic parameters were evaluated during the second saturating light period.

Under these two light regimes, photosynthetic performance was assessed using pulse amplitude modulated (PAM) fluorometry, measuring nonphotochemical chlorophyll fluorescence quenching (NPQ) as a proxy for the ability of algae to dissipate excess absorbed light energy into heat^71^. In addition, photosystem II (PSII) quantum yield (Φ_PSII_) and maximal PSII quantum yield (F_v_/F_m_) were measured as indicators of photosynthetic efficiency^72^.

Cultures subjected to the fluctuating light regime displayed lower quantum yields indicative of photoinhibition^73^, and decreased NPQ development, when compared to cultures that experienced a single exposure (Figure 3A-C). The latter observation suggests that the majority of light-dissipating capacity had already been initiated during the first saturating light period. Under these fluctuating conditions, addition of vitamin D significantly increased the quantum yields and NPQ observed during the second saturating light period (Figure 3A-C). These findings suggest that the vitamin D treatment facilitated a quicker relaxation of NPQ and PSII reaction centers after the initial exposure period to saturating light, thereby preparing the cell for subsequent exposure. Indeed, vitamin D treatment did not have a noticeable effect on algal cultures exposed to a single saturating-light period where relaxation of NPQ and PSII reaction centers was not required. These observations suggest that vitamin D plays a role in algal physiology under harmful light, particularly under fluctuating light levels.

**Figure 3.**
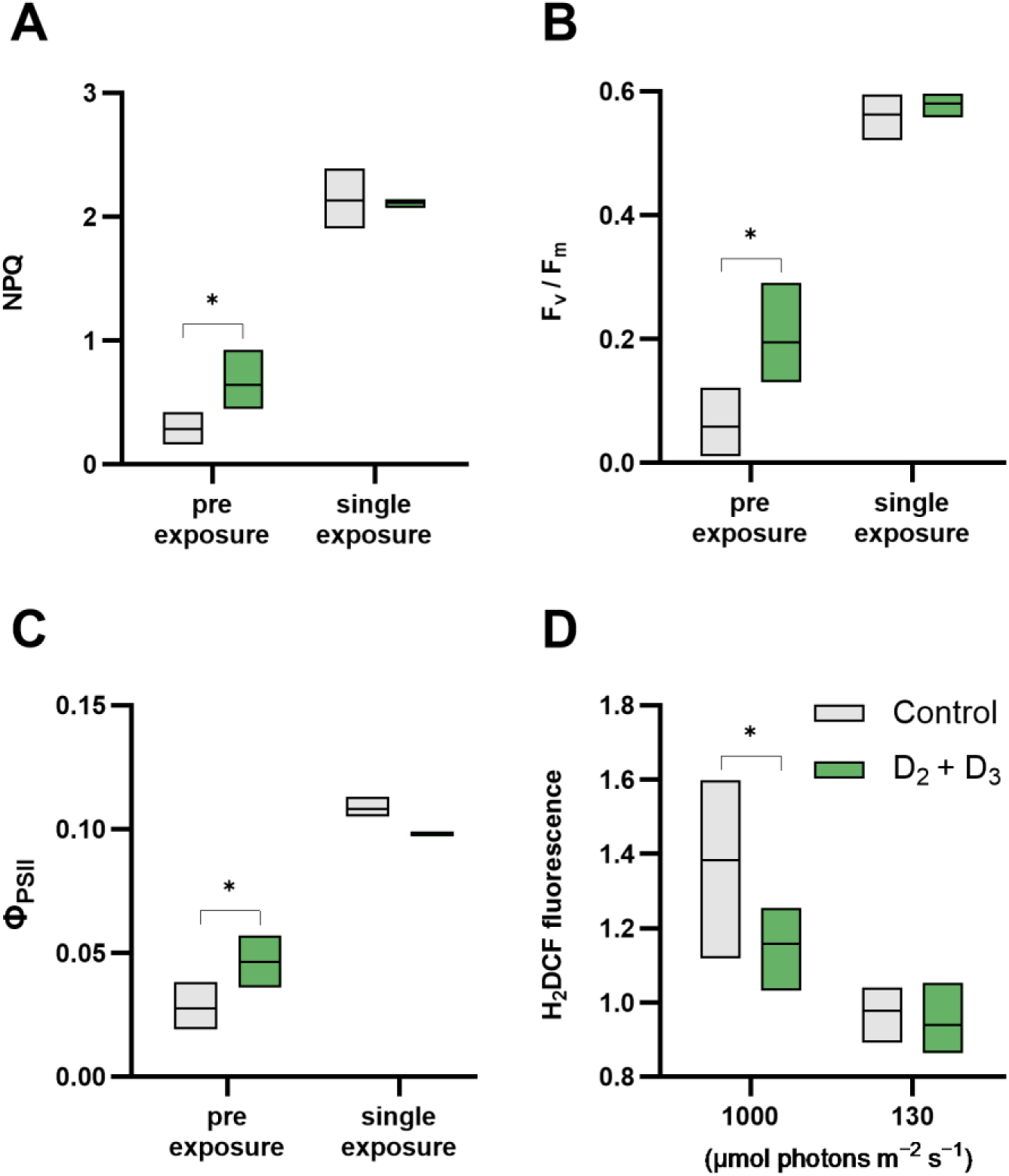
Vitamin D treatment improves the algal photosynthetic performance and alleviates oxidative stress following exposure to excess light. (A) Non photochemical quenching (NPQ), (B) F_v_/F_m_ values and (C) Φ_PSII_ values of vitamin D-treated and control algal cultures. Cultures were either pre-exposed to saturating light of 1000 μmol photons m^-2^ s^-1^ followed by dark incubation and a second exposure during the PAM analysis (“pre-exposure”), or not pre-exposed (“single exposure”). (D) Fluorescence values of vitamin D-treated and control algal cultures stained with the intracellular reactive oxygen species (ROS) probe H_2_DCF-DA. Cultures were exposed to regular light or excess light (130 or 1000 μmol photons m^-2^ s^-1^, respectively) intensities for 3 hours. Algae were cultivated in 50 ml and sampled for analysis at day 10 of growth. Statistically significant values (*p* < 0.05) of vitamin D-treated cultures compared to control conditions are marked by *, calculated based on three biological replicates using two-tailed paired t-test for (A-C) and one-tailed paired t-test for (D).

Interestingly, previous reports indicated that UV irradiation led to an increase in NPQ in *E. huxleyi*^58,74^. However, in our experiments, cultures subjected to UV did not exhibit a comparable effect (Figure S5).

### Vitamin D alleviates ROS accumulation under saturating light

Under conditions of saturating light, photosynthetic organisms are likely to experience photooxidative stress^75^, resulting in the production of detrimental ROS. Our qRT-PCR analysis detected the upregulation of two ROS mitigating mechanisms following treatment with vitamin D (Figure 2). One mechanism involves the LHCSR protein and operates by reducing ROS formation in the light-harvesting complex under excess-light by means of NPQ (G18590), and the other mechanism involves the enzyme heme oxygenase that produces antioxidants (G2511).

To further explore the possible role of vitamin D in regulating cellular ROS levels, we examined whether the addition of vitamin D to algal cultures under saturating light leads to reduced ROS levels in algae. We therefore subjected algal cultures to saturating light (1000 μmol photons m^−2^ s^−1^) and assessed the formation of intracellular ROS using the cell-permeable fluorescent probe 2,7-Dichlorodihydrofluorescein diacetate (H_2_DCF-DA). The tested algal cultures were either treated with D_2_ + D_3_ (0.5 µM of each) or supplemented with DMSO (untreated). Additionally, control cultures were exposed to regular light intensities (130 μmol photons m^−2^ s^−1^) and subjected to the same procedures. Our findings revealed a significant decrease in intracellular ROS formation in algae that were treated with vitamin D and exposed to saturating light, compared with untreated algae under the same light regime (Figure 3D). The reduction in ROS formation was detected after 3 hours of incubation under saturating light. These findings provide additional support for the involvement of vitamin D in the algal response to high light conditions, underscoring its role in alleviating oxidative stress induced by ROS.

## Discussion

### Vitamin D alleviates algal photo-oxidative stress

The current study reveals a photoprotective role of vitamin D in a globally abundant marine microalgal species. Our findings suggest that the photochemical transformation of vitamin D serves as an indicator of exposure to harmful radiation, consequently enhancing the algal response to excess-light stress. This enhanced response is manifested by a quicker relaxation of NPQ mechanisms and PSII reaction centers under fluctuating light, along with an overall reduction in cellular ROS. Our data suggest that a significant aspect of the response triggered by vitamin D is aimed at mitigating photo-oxidative stress.

### Vitamin D may serve as an algal light indicator

Vitamin D has the potential to serve as a sensitive light indicator in phytoplankton, particularly under dynamic light environments. The ocean surface mixed layer can extend depths of over 200 meters^76^, and phytoplankton cells are transported vertically in this layer^77^. This vertical transport can lead to rapid changes in light intensity within a matter of hours^33,78^. In such a dynamic scenario, as cells ascend, they experience an increase in UV-B and subsequently may generate vitamin D. The specificity of the photochemical conversion of vitamin D under UV-B, coupled with its relatively high reactivity to these wavelengths^79^, suggests that it could function as a sensitive proxy for assessing exposure to UV-B radiation, high light levels, or fluctuations in light intensity.

### Limitations of exogenous vitamin D addition

Exogenous administration of vitamin D is common in mammalian cell research, typically by dissolving it in an organic solvent and adding the mixture into the supernatant of cell cultures^53,55,56^. Mammalian cells regularly absorb vitamin D from the extracellular matrix and harbor vitamin D receptors in their plasma membrane^80^. The extent of exogenous vitamin D absorption by *E. huxleyi*, its stability inside the cell, and the specific membranes or organelles it may permeate, are currently unknown. Despite these challenges, our observations reveal a measurable response of *E. huxleyi* to exogenous vitamin D, leading to a beneficial impact such as reduced ROS under saturating-light stress (Figure 3D). Furthermore, there appears to be a synergistic effect of combined D_2_ and D_3_ on growth and gene expression, which merits further exploration. Furthermore, among the genes displaying overexpression under both UV and vitamin D treatments, a distinctly greater differential expression was evident in cultures subjected to UV (Figure 2: G18590, G2511 and G25467). The stronger response to UV, as opposed to vitamin D, might be due to the exogenous administration procedure. Though, these differences might stem from a broader cellular response to UV radiation, triggering additional regulatory mechanisms that are not specifically responsive to vitamin D alone.

### UV is an important environmental factor impacting algal physiology

When studying algal physiology, the significance of UV as an influential environmental factor should be acknowledged. In experimental setups aimed at studying algal physiology and ecology, UV radiation has traditionally been excluded due to its detrimental effects. However, in the environment, algae regularly encounter low levels of UV. While the omission of UV simplifies experimental conditions, our study unveils the influence of vitamin D, a product of UV-B exposure, on the response of *E. huxleyi* to environmental stress. These findings point to the potentially advantageous role of UV for algae facing excess-light stress and underscore the significance of studying algal physiology under environmentally relevant conditions.

### Vitamin D in vertebrates versus algae

Our study offers comparative insights on the role of vitamin D in vertebrates and in *E. huxleyi* algae. The extensive knowledge on vitamin D biology primarily originates from research on humans and other vertebrates. Transposing this knowledge to *E. huxleyi* presents challenges due to significant phylogenetic and physiological differences. Nevertheless, parallels can be drawn. Vitamin D was shown to enhance cellular defense in human and mice keratinocytes against UV-induced oxidative stress and DNA damage^53,54^. Vitamin D has also been shown to mitigate oxidative stress in rat liver and intestine^50,81^, partly through the upregulation of heme oxygenase, a response mirrored in vitamin D-treated algal cells.

### The possible ancient origin of vitamin D

Vitamin D synthesis likely has ancient origins, given its presence across various lineages of eukaryotes^9–15^. Sterols are a defining feature of eukaryotes, and the enzymatic pathways leading to the production of ergosterol and 7-dehydrocholesterol, which are precursor molecules to D_2_ and D_3_ forms of vitamin D, may have existed in the last eukaryotic common ancestor (LECA)^83,84^. Previous studies suggested the ancient evolutionary origins of vitamin D^20,85^, but the understanding of its role beyond the animal kingdom remained limited.

Eukaryotes likely evolved near oxygenic photoautotrophs in sunlit ocean surfaces due to the absence of oxygen in the deep ocean during their emergence^86^. Given the antioxidant properties of vitamin D and its evolutionary ties to oxidative agents like UV-B and oxygen, it raises the question of whether vitamin D evolved in early eukaryotes to sense oxidative environments. Similar roles have been proposed for sterols, particularly cholesterol, in response to environmental oxygen levels^87,88^. Thus, the unique reactivity of vitamin D to solar radiation may have played a role in the early stages of life on Earth.

## Materials and methods

### Algal strain and growth conditions

The axenic algal strain of *E. huxleyi* CCMP3266 was purchased from the National Center for Marine Algae and Microbiota (Bigelow Laboratory for Ocean Sciences, Maine, USA). Algae were grown in artificial sea water according to Goyet and Poisson^89^ and supplemented with L1 medium according to Guillard and Hargraves^90^, with the exception that Na_2_SiO_3_ was omitted following the cultivation recommendations for this strain. Algae were grown in standing cultures in borosilicate Erlenmeyer flasks with an initial inoculum of 330 cells/ml, placed in a growth chamber at 18°C under a light/dark cycle of 16/8 hr. Growth light intensity during the light period was 130 µmoles photons m^−2^ s^−1^. Cultures used for metabolomics, transcriptomic, qPCR, PAM and ROS accumulation analysis were grown in 50 ml medium inside 250 ml borosilicate Erlenmeyer flasks. Cultures used for growth analysis under vitamin D treatments, ImageStream flow cytometry and chlorophyll measurements were grown in 20 ml medium inside 100 ml borosilicate Erlenmeyer flasks.

Diurnal UV irradiation during the light period was achieved by placing a UV-emitting light source (Exo Terra Reptile UVB150 25W, Hagen, Montreal, Canada) inside the algal growth chamber, at a distance of 20 cm from the culturing flasks. Algal cultures experienced UV-A intensity of 0.5 w/m^2^, UV-B intensity of 0.07 w/m^2^ and UV-C intensity of 0.026 w/m^2^, measured using an ALMEMO 2470 data logger equipped with FLA 623 UV-A (310 to 400 nm, maximum sensitivity at 355), UV-B (265 to 315 nm, maximum at 297), and UV-C (220 to 280 nm, maximum at 265) probes (Ahlborn, Budapest, Hungary). UV radiation intensity was measured from within the Erlenmeyer flask in dry conditions, therefore the reported intensities take into account attenuation by the borosilicate boundary. The UV-B intensity used in this study was selected to emulate environmental UV-B intensities encountered at the ocean surface in some locations^29^. The UV light-source was operating daily for 14 hours in parallel to the light period, starting one hour after illumination started, and ending one hour before illumination ended. This irradiation regime aimed to mimic a simplified day cycle including dawn and dusk periods.

Vitamin D treatment was conducted by dissolving vitamin D_2_ or D_3_ (Sigma-Aldrich, Burlington, Massachusetts, USA) in DMSO. The final DMSO concentration in cultures, including controls, was 0.1%.

Algal growth was monitored by a CellStream CS-100496 flow cytometer (Merck, Darmstadt, Germany) using a 561 nm laser and plotting the chlorophyll fluorescence at 702/87 nm against forward scatter.

### Vitamin D analysis

Metabolic analysis was conducted following Oberson^82^. Standards for D_2_, D_3_, ergosterol and 7-dehydrocholesterol were purchased in dry (Sigma-Aldrich) and dissolved in CHCl_3_. Standard solutions of the different metabolites were combined into a single solution and diluted to create a standard curve. All final standards and samples were spiked with 50 ng of vitamin D_2_-d_3_ (IsoSciences, Ambler, Pennsylvania, USA) serving as an internal standard. Algal samples were centrifuged, lyophilized and stored in -80°C until analysis. Saponification was achieved by resuspending samples in 108 µl 55% KOH, 192 µl ethanol, and 60 µl of 9% NaCl and 7.4% ascorbic acid, followed by homogenization and stirring at room temperature for 18 hours. Samples were then supplemented with 40 µl 10% NaCl and 300 µl of 20% ethyl acetate in heptane, vortexed extensively and centrifuged for 30 minutes. The upper phase was collected, and the process was repeated twice. Samples were evaporated, dissolved in 200 µl of 0.5% isopropanol in hexane and sonicated. Strata SI-silica 55 μm 70 A columns (Phenomenex, Torrance, California, USA) were used for solid phase extraction and were pre-conditioned with 1 ml of 50% CHCl_3_ in isopropanol, followed by two washes with 1 ml of hexane. Samples were then loaded onto the columns and washed with 0.5 ml of 0.5% isopropanol in hexane which were discarded and washed again with 2.5 ml of 2.5% isopropanol in hexane which were collected. Samples were evaporated and dissolved in 200 µl of PTAD in acetonitrile, sonicated, stirred at room temperature for 2 hours, centrifuged for 10 minutes and transferred into LC-MS vials. Samples were protected from light during the extraction process.

Due to the inconsistency in identification of D_3_ in algal samples, several technical adaptations regarding algal growth and sample collection were implemented and evaluated. To examine whether inconsistencies arise due to rapid D_3_ enzymatic degradation, algal cultures were immediately placed on ice, centrifuged in a cooled, 4°C centrifuge, and the supernatant quickly discarded and replaced with 50% methanol in DDW. The samples were then plunged into liquid nitrogen and stored in -80°C. Later, samples were thawed, evaporated in vacuum to remove the methanol, lyophilized and proceeded to vitamin D extraction. Additional modifications included increasing the intensity of UV radiation during algal growth, increasing sample size by combining separate cultures, and using F/2 trace metal mix instead of L1 trace metals. These attempts did not improve the reproducibility of D_3_ detection.

Vitamin D was measured using an UPC2-ESI-MS/MS equipped with Acquity UPC2 system (Waters, Milford, Massachusetts, USA). The MS detector (Waters TQ-XS) was equipped with an ESI source. The measurements were performed in the positive ionization mode using MRM. The source and de-solvation temperatures were maintained at 150°C and 500°C, respectively. The capillary voltage was set to 1.5 kV. Nitrogen was used as the de-solvation gas and cone gas at a flow rate of 700 L h^-1^ and 150 L h^-1^, respectively. Ionization parameters of ergosterol, 7-dehydrocholesterol, D_2_ and D_3_ were adjusted by direct infusion of standards. Ionization parameters for other compounds were taken from Oberson^82^.

UPC2 system: mobile phase A consisted of CO_2_, and mobile phase B consisted of 98% MeOH, 2% DDW and 10mM ammonium formate. Make up solvent was 1% formic acid in 90% MeOH and 10% DDW at a flow rate of 0.4 ml min^-1^. The column (WATERS Acquity CSH FluoroPhenyl 1.7 µm, 3.0x100 mm, cat. 186006573) was maintained at 45°C, injection volume was 3 µl. At the first 0.5 min of injection, 99.5% of mobile phase B, and 0.5% of mobile phase A were run at flow rate of 2.0 ml min^-1^. Then, mobile phase A was gradually reduced to 92% at 6 min, and further decreased to 70% at a flow rate of 1.75 ml min^-1^ at 6.5 min. This composition of mobile phase and flow rate were kept until 7 min, followed by increase in mobile phase A to 99.5% at 7.8 min, and then increase in flow rate to 2 ml min^-1^ at 8.5 min, and running at those conditions until 9 min.

### RNA extraction

Algal cultures were harvested for RNA extraction by centrifugation at 4000 rpm for 5 min at 18°C. RNA was extracted using the Isolate II RNA mini kit (Meridian Bioscience, London, UK) according to manufacturer instructions. Cells were ruptured in RLT buffer containing 1% β-mercapto-ethanol by bead beating for 5 min at 30 mHz. RNA was then treated with 3 µl Turbo DNAse (ThermoFisher, Waltham, MA, USA) in a 50 µl reaction volume, followed by a cleaning step using RNA Clean & Concentrator^TM^-5 kit (Zymo Research, Irvine, CA, USA) according to manufacturer instructions. RNA yields were ∼200 µg/µl or higher, and RNA integrity was assessed by TapeStation 4150 (Agilent, Santa Clara, CA, USA). RNA was used for generating transcriptomic data and qRT-PCR analysis.

### Transcriptomic analysis

Transcriptomic data was generated using the MARS-seq library preparation protocol^91^, and analyzed with the UTAP pipeline^92^. As part of the pipeline, read counts for each gene were normalized using the DESeq2’s median of ratios method^93^. Differential expression (DE) between treatments was calculated using the following thresholds: mean number of normalized reads across all samples ≥ 5, adjusted p-value ≤ 0.05, Log2 fold change ≤ -0.7 or ≥ 0.7. The previously generated *E. huxleyi* CCMP3226 synthetic genome (sGenome) and annotation file was used as reference for the UTAP pipeline^94^. Briefly, the *E. huxleyi* CCMP3226 sGenome was generated by *de novo* transcriptome assembly of short-reads and long-reads. The assembled *E. huxleyi* CCMP3226 transcripts were then mapped to the *E. huxleyi* CCMP1516 reference genome^95^ to define gene loci. For the current work, functional gene annotations were manually curated by identifying open reading frames in assembled transcripts using the ORF finder tool (www.ncbi.nlm.nih.gov/orffinder; transcript accessions are given in Table S1) and analyzing protein domains in the translated sequences using InterProScan 5^96^. Additionally, transcript sequences were searched against the SwissProt database using NCBI blastx^97^, and the hit with the highest E-value taken. Specifically, the gene loci analyzed using blastx were G18590, sharing highest similarity to *Chlamydomonas reinhardtii* LHCSR (NCBI accession P93664.1) with E-value of 1e-26 and nucleotide identity of 57%; G25467, sharing highest similarity to rat *cyp1a2* (NCBI accession P04799.2) with E-value of 1e-31 and nucleotide identity of 27%; G14502, sharing highest similarity to *Arabidopsis thaliana* Calmodulin-like protein 12 (NCBI accession P25071.3) with E-value of 6-e5 and nucleotide identity of 22.6%. The putative Ca-binding activity of G26534 was assessed by identifying *bona fide* Ca-binding domains using InterProScan 5. Specifically, we performed blastx^97^ and focused on the highest hit that contained an identifiable protein domain using InterProScan 5^96^, resulting in the identification of an EF-hand family protein in *Chrysochromulina tobinii* that harbors three EF-hand domain pairs (NCBI accession KOO34173.1, with E-value of 8e-13 and nucleotide identity of 32%).

### Imaging flow cytometry and pigment analysis

For imaging flow cytometry, cultures were grown in 20 ml medium and either exposed to UV daily or treated with vitamin D_2_ and D_3_ (0.5 µM of each) at day 4 of growth. Control and UV-exposed cultures were treated with equal amounts of DMSO. Cultures were analyzed by imaging flow cytometry (ImageStreamX, Amnis, Cytek Biosciences, Fremont, California, USA). Side scatter was determined using 785 nm laser (3.75mW), chlorophyll fluorescence using a 405 nm laser with collection at 640-745 nm (channel 11), and brightfield using channels 1 and 9 of the device. At least 2600 cells were collected from each sample. Data were analyzed using image analysis software (IDEAS 6.3; Amnis). Cells were gated for single cells using the area and aspect-ratio features on the brightfield channel, and for focused cells using the gradient root mean square (Gradient RMS)^98^ and Contrast features. Cells were further gated for chlorophyll positive using the area and intensity of the chlorophyll channel. To calculate cell area, the Object mask was used (delineates the cell morphology) on the brightfield image. To calculate chlorophyll area, the Morphology mask was created on the chlorophyll channel. The number of chloroplasts was calculated using the Spot count feature on the Peak mask (value set at 1.1), on the chlorophyll channel.

Chlorophyll a was measured by filtering at least ∼4 * 10^6^ cells from each culture on a Whatmann GF/C filter and extraction with 3 ml methanol overnight at 4 °C. Samples where centrifuged for 10 minutes, 1 ml of extract was placed in a polystyrene cuvette and measured by Ultrospec 2100 pro (Biochrom, Cambridge, UK). Chlorophyll a concentration as µg/ml was measured by [Chl a] = (16.29 * E^665^) – (8.54 * E^652^), where E^665^ and E^652^ are the absorption at 665 and 652 nm, respectively, after deduction of the absorption at 750 nm^99^. The concentration per ml was further multiplied by the extraction volume and divided by the total number of cells filtered.

### Quantitative real time PCR (qRT-PCR)

Algal cultures were treated with 1 µM of vitamin D species as described earlier. UV-treated cultures were exposed to the same UV intensities as described previously and treated with equal amounts of DMSO. All treatments lasted 1 hour. Equal concentrations of RNA taken from 10 days old cultures were utilized for cDNA synthesis using Superscript IV (ThermoFisher), according to manufacturer instructions. qPCR was conducted in 384 well plates using SensiFAST SYBR Lo-ROX Kit (Meridian Bioscience, Cincinnati, OH, USA) in a QuantStudio 5 qPCR cycler (Applied Biosystems, Foster City, CA, USA). The qPCR program ran according to enzyme requirements for 40 cycles. Samples were normalized using three housekeeping genes: *alpha-tubulin*, *beta-tubulin* and *ribosomal protein l13 (rpl13*). DNA contamination was assessed by applying the same program on RNA samples that were not reverse transcribed (omitting the Superscript IV enzyme in the reverse transcription reaction mix). Gene expression ratios were analyzed according to Vandesompele^100^ by geometric averaging of housekeeping genes. Relative gene expression levels were compared to control samples. Primer efficiencies were determined using the QuantStudio 5 software, by qPCR amplification of serially diluted cDNA. All primers had a measured efficiency between 80-120%. Primer sequences are given in Table S2.

### Pulse amplitude-modulated fluorometry (PAM) analyses

For PAM analyses, algal cultures at day 10 of growth were divided into four subcultures (two pairs) that were subjected to different treatments. Under the ‘single exposure’ assay, one subculture was treated with D_2_ + D_3_ (0.5 µM of each) and the second one treated with an equal amount of DMSO (control). Following 2 hours, the two subcultures were incubated in the dark for 5 minutes and analyzed using WATER-PAM II (Heinz Walz GmbH, Effeltrich, Germany), applying saturating pulses of 0.9 s, 6000 µmol photons m^−2^ s^−1^, and actinic light intensity of 1150 µmol photons m^−2^ s^−1^. Under the ‘pre exposure’ assay, the second pair of subcultures was subjected to the same vitamin D and control treatments, and placed in the growth chamber under saturating light intensities of 1000 µmol photons m^−2^ s^−1^ using white LED lamps. Following 2 hours of vitamin D and saturating light treatments, the paired subcultures were incubated in the dark for 5 minutes and analyzed as described above.

Maximum PSII quantum yield (F_v_/F_m_) was calculated as F_v_/F_m_ = (F_m_ - F_0_) / F_m_, where F_o_ is the baseline fluorescence under a measuring light of 160 μmol photons m^−2^ s^−1^ and F_m_ is the maximum fluorescence yield at the first saturating pulse^101^. PSII quantum yield (Φ_PSII_) was calculated as (F’_m_ –F_t_) / F’_m_, where F’_m_ is the maximum fluorescence yield at a given saturating pulse, and F_t_ the steady-state fluorescence value under actinic light immediately prior to the saturating pulse. Non-photochemical quenching (NPQ) was calculated as NPQ = (F_m_ –F’_m_) / F’_m_^101^.

### Intracellular reactive oxygen species (ROS) measurements

Algal cultures were divided into four subcultures (two pairs) and treated with D_2_ + D_3_ (0.5 µM of each) or DMSO as described under ‘chlorophyll fluorescence’. Following 2 hours of vitamin D or DMSO treatment, the subcultures were stained with 0.5 μM of H_2_DCFDA (ThermoFisher) in DMSO and incubated in the dark for 20 minutes. Then, one pair was returned to regular light conditions of 130 µmol photons m^−2^ s^−1^ and the second pair exposed to saturating light of 1000 µmol photons m^−2^ s^−1^ using white LED lamp for 3 hours. DCF fluorescence was measured using CellStream CS-100496, excited at 488 nm and the signal was collected at 528/46 nm. The algal population was gated by plotting chlorophyll fluorescence (excitation-emission 561-702/87 nm) against forward scatter.

## Data Availability

The transcriptomics data discussed in this publication have been deposited in NCBI’s Gene Expression Omnibus and are accessible through GEO Series accession number GSE243677 (https://www.ncbi.nlm.nih.gov/geo/query/acc.cgi?acc=GSE243677)

## Acknowledgments

We appreciate the technical guidance of Dr. Shifra Ben-Dor, Dr. Merav Kedmi and Dr. Hadas Keren-Shaul in RNA-sequencing and are thankful for the help of Dr. Ron Rotkopf with statistical analysis, and of Dr. Alexander Brandis with LC-MS analysis (Life Sciences Core Facilities, Weizmann Institute of Science, Israel). We thank Dr. Shilo Rosenwasser (The Hebrew University of Jerusalem, Israel) for sharing his expertise in pulse amplitude-modulated fluorometry. We thank Prof. Robert Fluhr and Prof. Dan Yakir (Weizmann Institute of Science, Israel) for valuable comments during the study. We are grateful for Dr. Sheera Adar and Yuval Cohen (The Hebrew University of Jerusalem, Israel) for their insights into UV-induced DNA damage, and to Dr. Chana Kranzler (Bar Ilan University, Israel) for her technical and scientific support. Finally, we thank all members of the Segev lab for insightful discussions and input. O.E. received the Sustainability and Energy Research Initiative (SAERI) fellowship. The study was supported by funds received from the Weizmann SAERI program, the European Research Council (ERC StG 101075514), the Israeli Science Foundation (ISF 947/18), and the de Botton Center for Marine Science, granted to E.S.

## Authors Contributions

O.E. and E.S. designed the study. O.E., S.M. and I.P. performed and analyzed experiments. O.E., E. F., M.S. and Z. P. performed computational analyses. O.E. and E.S. wrote the manuscript. All authors discussed the results and contributed to the final manuscript.

