## Supplemental Information for "The Role of Vitamin D in *Emiliania huxleyi*: A Microalgal Perspective on UV Exposure"

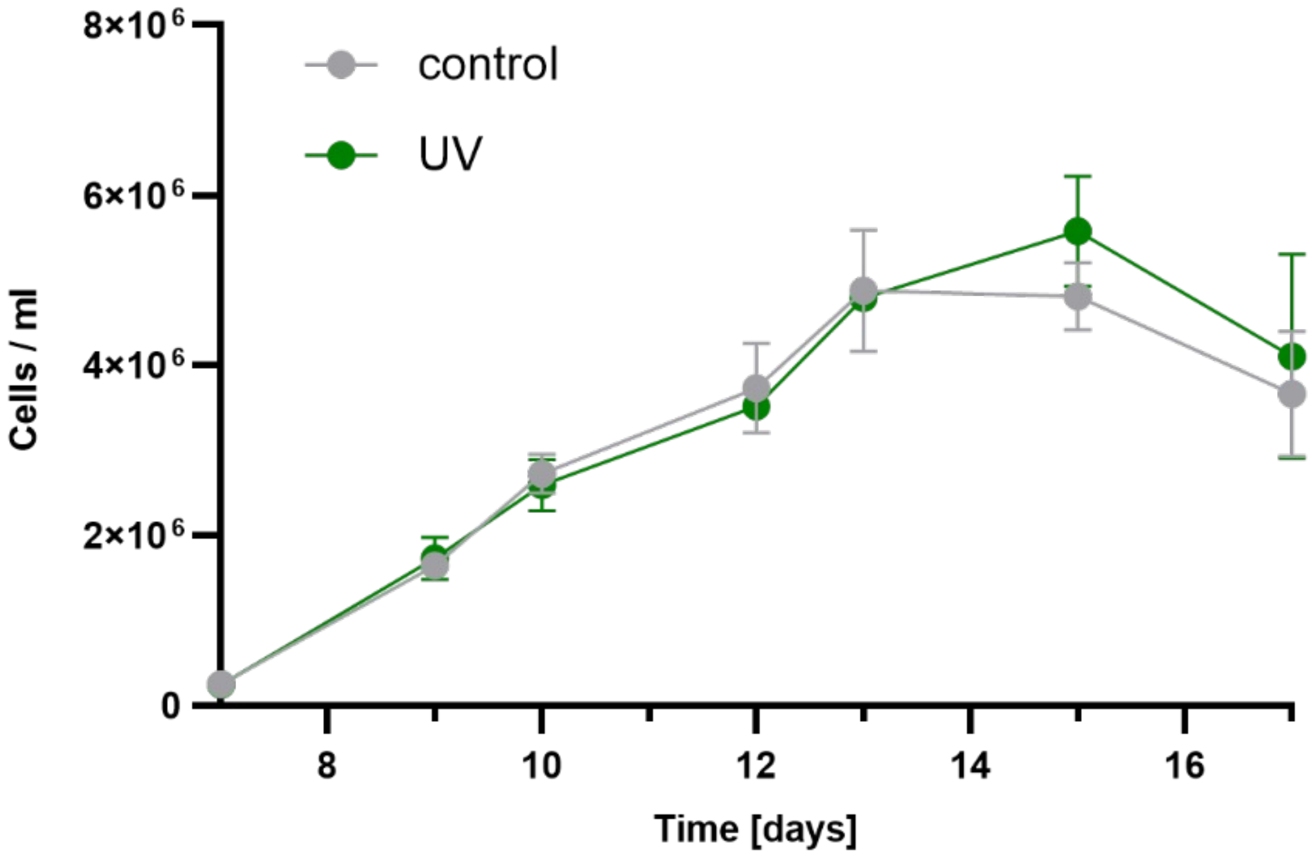


**Figure S1. Growth curves of algal cultures under control conditions and UV irradiation**. Algae were cultivated in 50 ml of growth medium, and RNA was harvested for RNAseq and qRT-PCR analyses. Cell density was measured from day 7. Error bars indicate standard deviation based on 3 biological replicates.

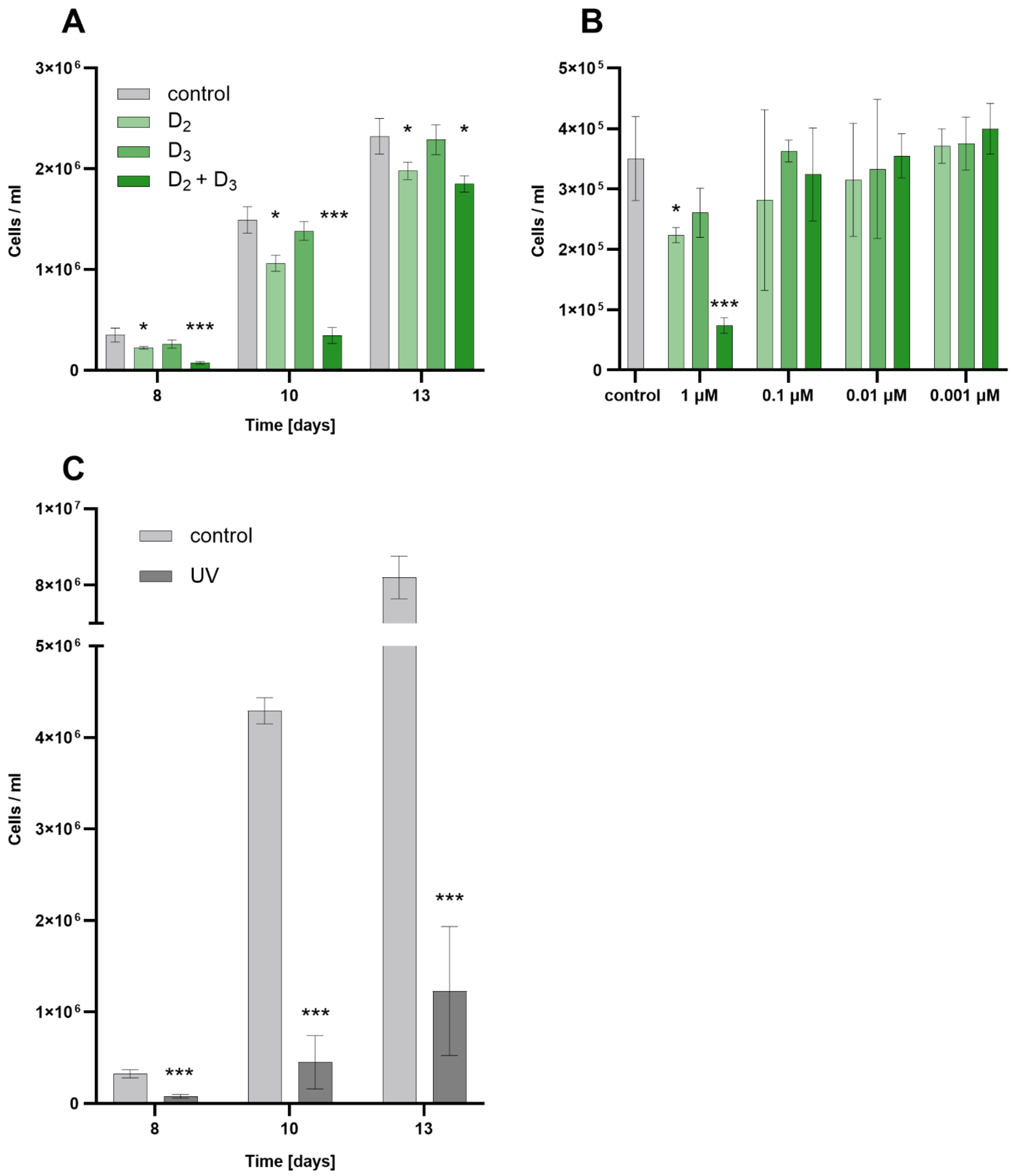


**Figure S2. Algal growth under vitamin D and UV.** (A) growth dynamics of algal cultures treated with 1μM of vitamin D species dissolved in DMSO: $D_2$, $D_3$, and both $D_2 + D_3$(0.5 μM of each). (B) Variation in algal cell densities at day 8 of growth under treatment with different concentrations of vitamin species. (A,B) share the same legend. Vitamin D was supplemented to cultures once at day 4. Control cultures were supplemented with identical volumes of DMSO. (C) growth dynamics of algae cultivated in 20 ml growth medium under daily UV irradiation compared to control conditions. Error bars indicate standard deviation based on 3 biological replicates. Statistical significance of treated cultures compared to control conditions of the same time point was calculated using two-tailed t-test assuming equal variances. One, two or three asterisks indicate $p < 0.05$, $p < 0.01$ and $p < 0.001$, respectively.

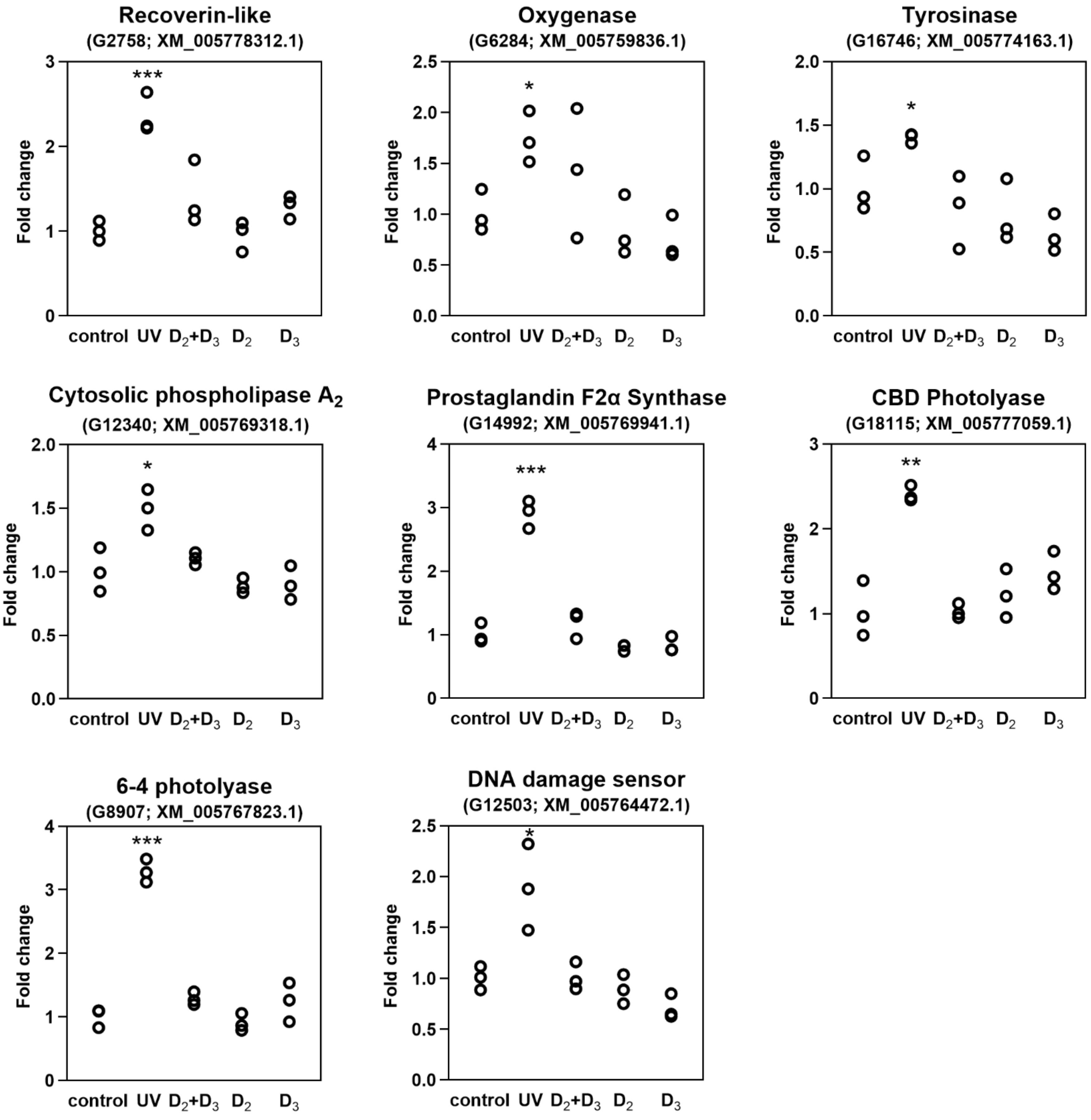


**Figure S3. UV irradiation results in upregulation of various algal signaling and stress-response mechanisms.** qRT-PCR analysis of genes under 1 hour of UV exposure or vitamin D treatments. Top title denotes gene products. In brackets: gene identifier in *E. huxleyi* CCMP3266[1] and matching gene in *E. huxleyi* CCMP1516 reference genome[2]. Statistical significance compared to control was calculated using two-tailed t-test assuming equal variances. One, two or three asterisks indicate $p < 0.05$, $p < 0.01$ and $p < 0.001$, respectively.

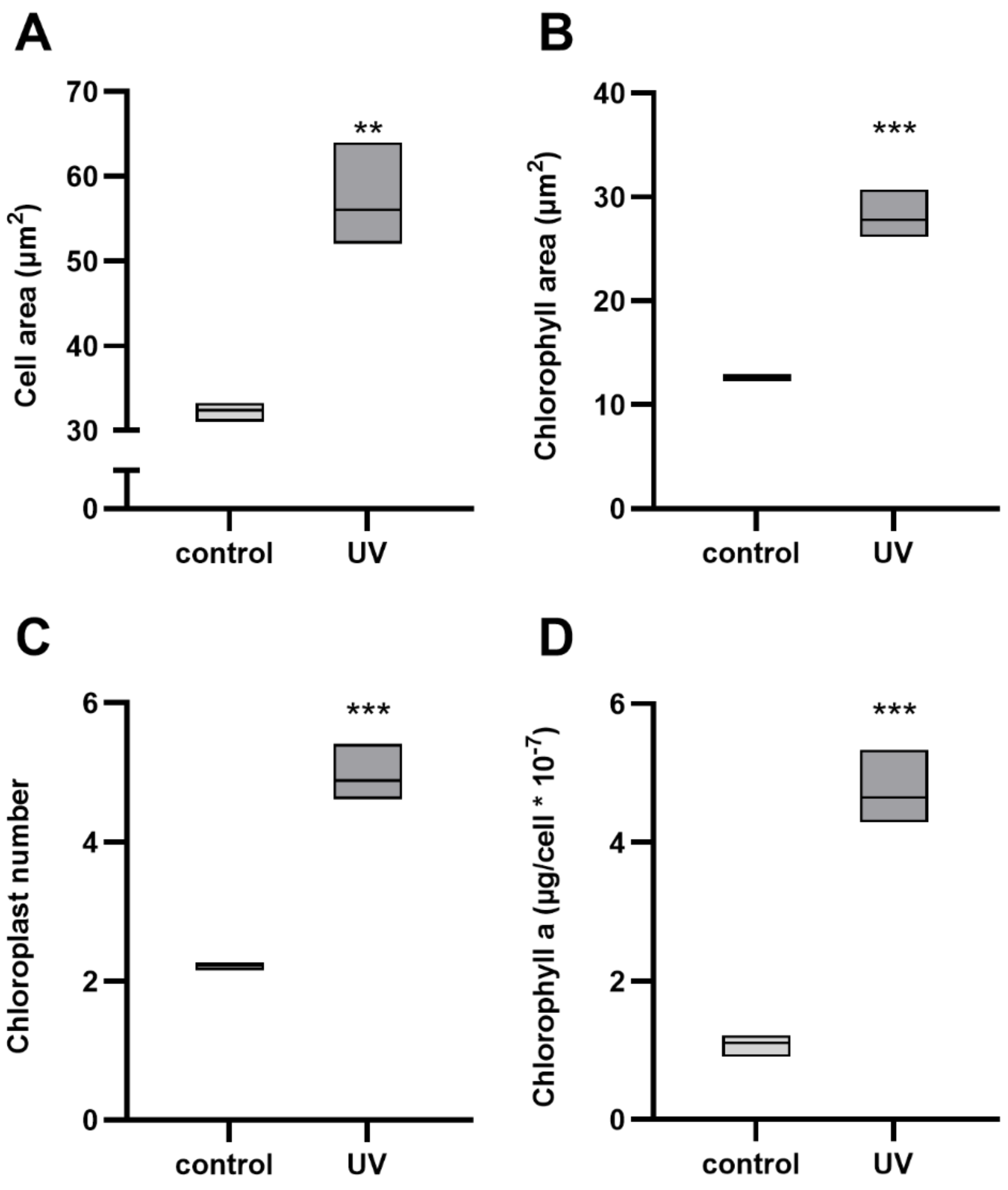


**Figure S4. Growth under UV irradiation leads to increased cell size, chloroplast size, chloroplast number and cellular chlorophyll a content.** (A) cell area, (B) chlorophyll area and (C) average number of chloroplasts analyzed using imaging flow cytometry, and (D) cellular chlorophyll a content. Cultures were either grown under daily UV irradiation or protected from UV as control. Statistical significance of treated cultures compared to control conditions was calculated based on three biological replicates for (A-C) and six biological replicates for (D) using two-tailed t-test assuming equal variances. Two or three asterisks indicate $p < 0.01$ and $p < 0.001$, respectively.

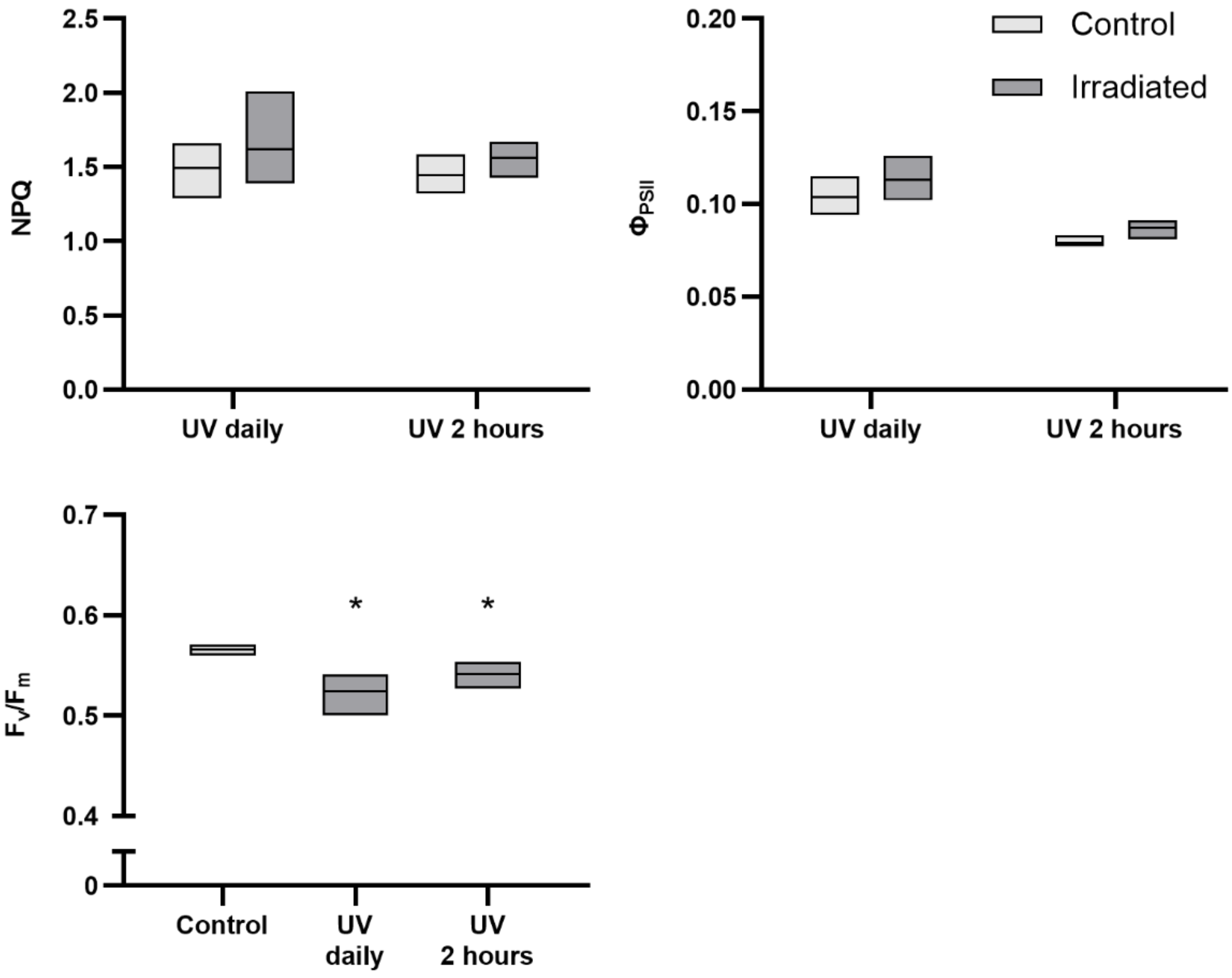


**Figure S5. UV radiation does not affect NPQ and $\Phi_{PSII}$ in *E. huxleyi*.** Cultures were either exposed to UV daily during the light period, once for 2 hours, or not exposed (control), and incubated in the dark for 5 minutes prior to PAM analysis. Analysis was conducted using cultures grown in 20 ml, at day 10 of growth. Statistical significance ($p < 0.05$) compared to control is marked by *, calculated from three biological replicates using two-tailed t-test assuming equal variances.

| Gene Identifier | Forward | Reverse |
|---|---|---|
| G6284 | GCCCTACCGGGTGTTTATCC | CTCGACTTGGTTGAGAACTTGC |
| G18115 | CACCGAGCCGACCCAAAAT | TTCATCTCGGTCGTTGACAGG |
| G18590 | CTCCTCGCGATGCAGAACAA | CCCTTTCTGCCAACGTGATCT |
| G25467 | TACGAGAATCGGCTGCTACG | CCCGTCGGACCTTAAGACAG |
| G12503 | TCTCGGTGGAAATGGCGAC | ATTGTCATCGAGGCCGCAAA |
| G93 | GTCACGCCGGCGACAAA | GCGATGTGCGGGTGTATCT |
| G12340 | AACCTGCTTGCCGACATGAT | GGTTGAATCAGCATTGACCCC |
| G14992 | GCGGGCTCTACTGAATCCG | GGGTCCTCGTAGAAGGTGTG |
| G16746 | TCGAAGATCCGGACGACGAT | AGCGCGAGACGAAAATAACG |
| G2511 | TTGTCGCGTCGCTCTACTTT | GCCGAAGTAGTACGCCATGT |
| G2758 | ATGGACCTAGACTCGGACGG | ACAGCCCTCAAGCTCACATC |
| G8907 | CTCTCGTGCTCGTGCTTCTT | TCGTAGATGTACTTTGCCGGG |
| G26534 | CTGGAAGATCGAGGCAACGG | TATGGCGTCGCCGTCAAAG |
| G26797 alpha-tubulin | CGAGAAGGCGTACCACGAG | CTTCGTCTTGATGGTGGCGA |
| G28192 beta-tubulin | CAACATGAAGTGCGCCATCT | CCTCGGTGAACTCCATCTCG |
| G1895 rpl13 | ACCAGCACTTCCACAAGACG | TGCCGCAGCTTGTAGTTGTA |

**Table S2. Primers used in this study.** Gene identifiers denote genes in *E. huxleyi* CCMP3266[1]. Primers were used for qRT-PCR analyses.
